# O-specific antigen-dependent surface hydrophobicity mediates aggregate assembly type in *Pseudomonas aeruginosa*

**DOI:** 10.1101/2021.01.14.426723

**Authors:** Sheyda Azimi, Jacob Thomas, Sara E. Cleland, Jennifer E. Curtis, Joanna B. Goldberg, Stephen P. Diggle

## Abstract

Bacteria live in spatially organized aggregates during chronic infections, where they adapt to the host environment, evade immune responses and resist therapeutic interventions. Although it is known that environmental factors such as polymers influence bacterial aggregation, it is not clear how bacterial adaptation during chronic infection impacts the formation and spatial organization of aggregates in the presence of polymers. Here we show that in an *in vitro* model of cystic fibrosis (CF) containing the polymers eDNA and mucin, O-specific antigen is a major factor in determining the formation of two distinct aggregate assembly types of *Pseudomonas aeruginosa* due to alterations in cell surface hydrophobicity. Our findings suggest that during chronic infection, interplay between cell surface properties and polymers in the environment may influence the formation and structure of bacterial aggregates, which would shed new light on the fitness costs and benefits of O-antigen production in environments such as CF lungs.

**Importance:** During chronic infection, several factors contribute to the biogeography of microbial communities. Heterogeneous populations of *Pseudomonas aeruginosa* form aggregates in cystic fibrosis airways, however, the impact of this population heterogeneity on spatial organization and aggregate assembly is not well understood. In this study we found that changes in O-specific antigen determine the spatial organization of *P. aeruginosa* cells by altering the relative cell surface hydrophobicity. This finding suggests a role for O-antigen in regulating *P. aeruginosa* aggregate size and shape in cystic fibrosis airways.

## Introduction

During chronic infection, biofilm-forming cells are often more tolerant to antibiotics and the host immune response than planktonic cells (1-3). Biofilms also allow individual cells the physical proximity to engage in and benefit from social behaviors such as quorum sensing (QS) and the production of secreted common goods (4-9). Biofilms formed during infection often take the form of cellular aggregates (8, 10-12). In the fluids of wounds and airways of cystic fibrosis (CF) patients, the opportunistic pathogen *Pseudomonas aeruginosa* frequently grows as freely suspended aggregates of ∼10-10,000 cells (6, 8, 10, 11). The mechanisms that govern the shape and size of bacterial aggregates during infection are not well defined. In polymer rich environments, aggregates have been shown to form either by (i) an increase in entropic force (depletion aggregation) (13, 14); or (ii) electrostatic interactions between bacterial cell surfaces and polymers in the environment (bridging aggregation) (15-18).

*P. aeruginosa* populations become phenotypically and genetically diverse over time in the complex micro-environments found in CF airways (19-22), and the importance of this heterogeneity on the formation and organization of aggregates has not been assessed. To resolve this, we evaluated aggregate formation in seven genetically diverse isolates sourced from heterogeneous populations of the *P. aeruginosa* strain PAO1 previously evolved in biofilms for 50 days (23). We assessed how each isolate formed aggregates in a polymer-enriched, spatially-structured CF growth medium (SCFM2: containing mucin and eDNA polymers), which has previously been used as an *in vitro* CF model to study the biogeography and physiology of *P. aeruginosa* (8, 24-26).

We found that the PAO1 ancestor and five isolates formed a stacked pattern (stacked aggregates) where cells closely packed lengthwise, similar to those identified in previous polymer driven depletion-aggregation studies (27). Two isolates formed distinct disorganized aggregates of varying sizes (clumped aggregates), similar to aggregates previously observed in CF sputum samples (10, 28). Whole genome sequencing showed that the two clumping isolates had alterations in the *ssg* gene (PA5001), which has previously been shown to be involved in lipopolysaccharide (LPS) core and O-antigen biosynthesis (29-32). In *P. aeruginosa*, LPS contains three major components: the lipid A layer of the outer membrane, a core oligosaccharide and O-antigen components. O-antigens are further subdivided into a D-rhamnose homopolymer found in most strains called the common polysaccharide antigen (CPA, formally A-band) and a variable heteropolymer of 3 to 5 sugars called the O-specific antigen (OSA, formally B-band) that confers serotype specificity (29, 32).

We hypothesized that changes in O-antigen glycoforms capping LPS would lead to different aggregate assembly types by altering the physiochemical properties of the bacterial cell surface. This could result in new interactions (e.g. between the bacteria or the bacteria and polymers) that compete with the entropic force that otherwise leads to stacked aggregation in this polymer-rich environment. To elucidate the contribution of LPS-capping glycoforms to aggregate assembly, we assessed the aggregate formation of clean deletion mutants of OSA and CPA. We found that loss of OSA and changes in the capped LPS core glycoforms led to increased hydrophobicity of the cell surface that overcame the entropic force imposed by host polymers, resulting in disorganized, irreversible clumping. Most importantly, we demonstrate that aggregate shape and structure is dependent on the interplay between the physical properties of the environment and the biological mediation of bacterial cell surface properties governed by the LPS core and OSA.

## Results

### Distinct aggregate assembly types in genetically diverse *P. aeruginosa* isolates

Genetic and morphologically heterogeneous isolates of *P. aeruginosa* are commonly collected from expectorated CF sputum samples (20, 21, 33, 34). Since it is known that several lineages of *P. aeruginosa* can stably coexist in CF airways, we tested whether population heterogeneity impacted aggregate formation. We chose seven distinct morphotypes isolated from a previous study where we evolved biofilms of PAO1 in a synthetic polymer-free sputum media (SCFM) for 50 days (23, 35, 36) (Fig. S1A). We assessed aggregate formation in a spatially structured iteration of SCFM termed SCFM2 which contains mucin and eDNA polymers (24). We identified two distinct types of aggregate assembly, where PAO1 and five of the evolved isolates (A2, B8, B13, C25 and D4) were assembled into stacked aggregates, where cells were closely aligned side by side by entropic force, similar to previous reports (27) (Fig. 1A; Fig. S1B & C). In contrast, two of the evolved isolates (A9, B9) formed clumped aggregates that appeared as disorganized small and large groups of cells, similar to bridging aggregation (15) (Fig. 1A). We also investigated growth of A9 and B9 in SCFM (no addition of eDNA or mucin) and observed that both formed clumps even in a polymer free environment, while the other isolates did not form any aggregates (Fig. S2A).

**Fig. 1.**
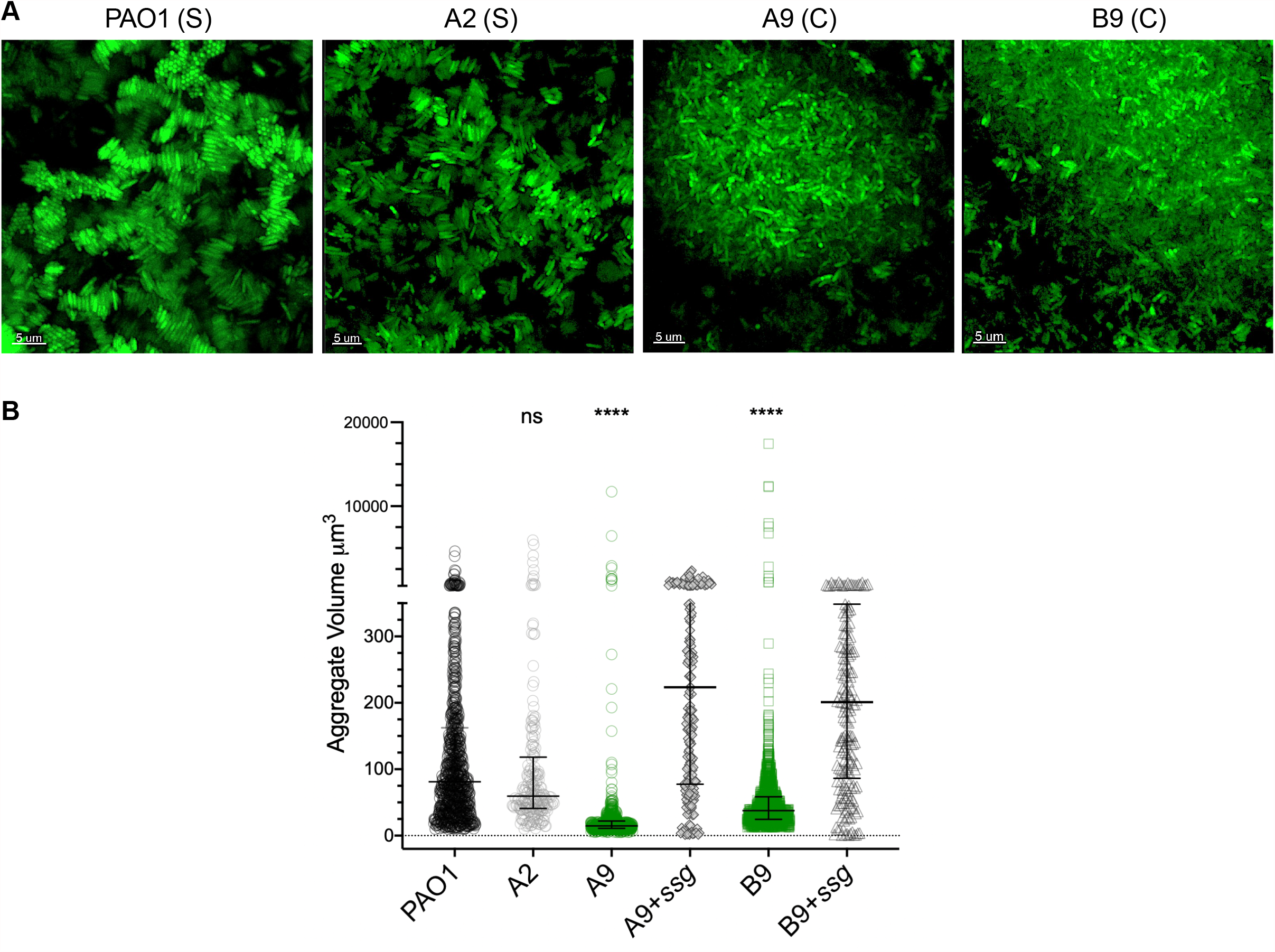
The two types of aggregate assembly formed by *P. aeruginosa* isolates in SCFM2 is due to *ssg* gene mutation. (A) In PAO1 and evolved isolates, aggregates assembled into either organized stacked structures (labeled as S) or disorganized clumps (labeled as C). (B) Stacked aggregates of PAO1 and A2 were significantly larger than aggregates formed by A9 and B9, and complementation with an intact *ssg* gene significantly increased the aggregate volume (Kruskal-Wallis, Dunn’s multiple comparison test, *p<*0.0001; error bars are median with interquartile range of aggregate volume, each data point is representative of an aggregate).

To identify genetic determinants of clumping assembly in the evolved isolates, we performed whole-genome sequencing on each isolate using the Illumina Miseq platform. We used breseq (0.34.0) for variant calling between the evolved isolates and the PAO1 ancestor (37). Whilst we found differential levels of genetic variation in each isolate when compared to PAO1, we observed that A9 and B9 each contained a 1 bp deletion in the *ssg* gene (PA5001) (Dataset 1). To confirm that mutation of *ssg* results in a clumped aggregate assembly, we complemented A9 and B9 with an intact *ssg* gene *in trans*. We found that in both isolates, *ssg* complementation restored the stacked aggregate assembly seen in PAO1 (Fig. 1B; Fig. S2B) suggesting that Ssg plays a role in aggregate assembly type.

A distinct feature of the stacked versus clumped aggregates was the average volume. We found that stacked aggregate volumes were 2-4 times larger (median of ≈ 63 µm^3^ and 89 µm^3^ aggregate size for PAO1 and A2 respectively) and (median of ≈ 200 µm^3^ in *ssg* complemented A9 and B9) compared to clumped aggregates (median aggregate size of ≈ 23 and 30 µm^3^ for A9 and B9 respectively) (Fig. 1B). To quantify these observed differences in the distribution of aggregate biomass in cells with stacked and clumped aggregate assembly types, we compared the distribution of biovolume (ratio of aggregate volume to surface area) for each type of aggregate assembly in SCFM2. We found that regardless of the size, the median biovolume in stacked aggregates was significantly higher than in clumped aggregates (Table S1).

### OSA and not other biofilm traits determines aggregate assembly type

The proposed function of Ssg is a glycosyltransferase, involved in LPS and exopolysaccharide biosynthesis (30, 38). *P. aeruginosa* with mutations in *ssg* have previously been shown to display decreased motility, enhanced phage resistance and a lack of O-antigen (30, 31, 39). We next determined whether the different aggregate assemblies (due to the loss of *ssg* in our evolved PAO1 isolates) was because of differences in O-antigen production. We constructed a clean *ssg* gene deletion in PAO1 (PAO1Δ*ssg*) and a range of isogenic LPS synthesis or O-antigen assembly mutants. These were (i) mannose reductase (PAO1Δ*rmd* OSA^+^, CPA^−^); (ii) epimerase (PAO1Δ*wbpM* OSA^−^, CPA^+^); (iii) OSA polymerase (PAO1Δ*wzy*, OSA^−^, CPA^+^); (iv) common initiating glycosyltransferase (PAO1Δ*wbpL*, OSA^−^, CPA^−^) and (v) O-antigen ligase (PAO1Δ*waaL*, OSA and CPA still made but not attached to LPS in the periplasm). In addition, we made mutants in the high (PAO1Δ*wzz1* OSA^+^, CPA^+^) and very high (PAO1Δ*wzz2*, OSA^+^, CPA^+^) OSA chain-length regulators (40, 41) (Fig. S3). We then determined the aggregate assembly type of the O-antigen mutants in SCFM2. We found that *ssg, wbpL* and *wbpM* mutants with no OSA, formed clumped aggregates, but lack of CPA alone (PAO1Δ*rmd*) did not change the aggregate assembly type (Fig. 2A & B).

**Fig. 2.**
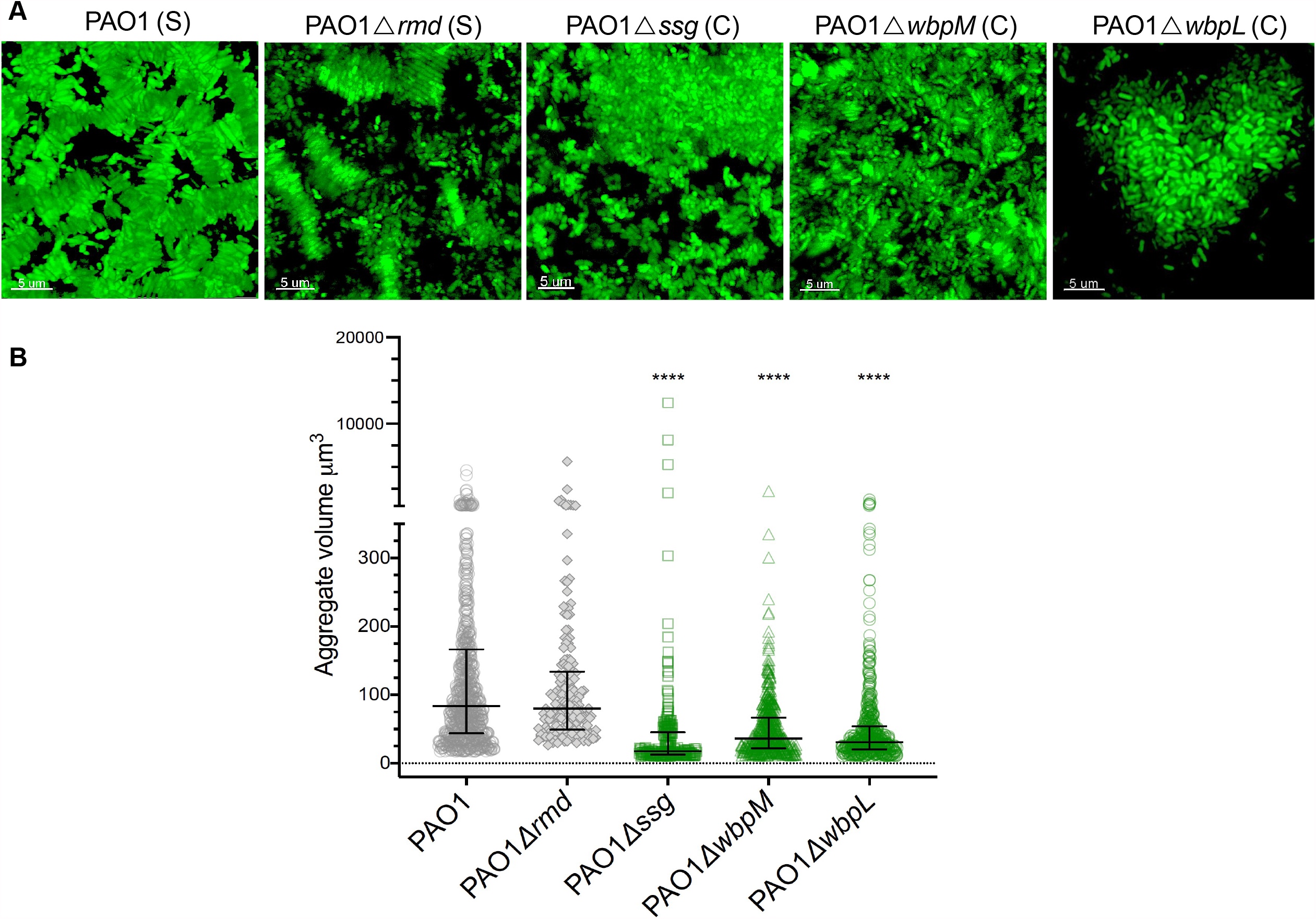
Loss of OSA leads to clumped aggregate assembly. (A) Loss of CPA (Δ*rmd*) did not alter the type of aggregate assembly, and loss of OSA (Δ*wbpM*) lead to dispersed small aggregates. The loss of both CPA and OSA(Δ*wbpL*) changed aggregate assembly type similar to the *ssg* mutant. (B) There was a significant reduction in aggregate volume in *ssg, wbpL* and *wbpM* mutants, however only the loss of *ssg* and *wbpL* displayed large, clumped aggregates (Kruskal-Wallis, Dunn’s multiple comparison test, *p*<0.0001; error bars are median with interquartile range of aggregate volume, each point is representative of an aggregate).

Biofilm formation by *P. aeruginosa* is regulated by several well-described mechanisms such as exopolysaccharide production, adhesins and quorum sensing (QS) (6, 9, 42, 43). To determine whether any of these factors interfered with stacked aggregation in SCFM2, we examined the aggregate assembly of defined mutants in exopolysaccharide production (PAO1Δ*pel/psl*), lectins (PAO1Δ*lecA &* PAO1Δ*lecB*) and a mutant lacking a major QS regulator (PAO1Δ*lasR*). Although the role of QS and biofilm formation remains controversial, we assessed this mutant because LasR regulates several pathways that could impact on aggregation type (7, 44). We found that all these mutants displayed stacked aggregate assembly like PAO1 (Fig. 3). This indicated that changes in aggregate assembly type was not due to alterations in common phenotypes associated with biofilm formation, only the loss of OSA was important.

**Fig. 3.**
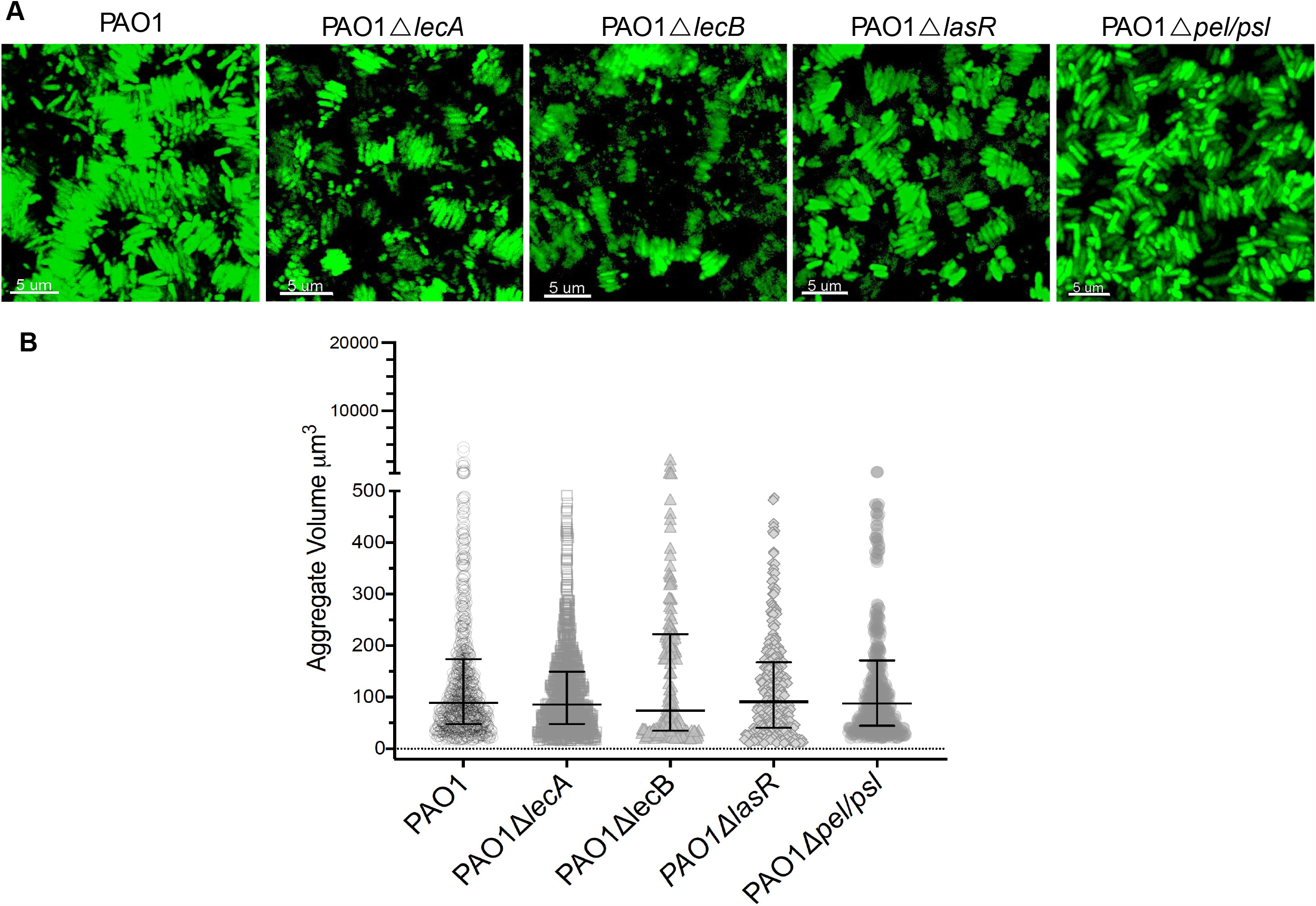
The aggregate assembly type is independent of exoploysaccharide production, lectins and quorum sensing. (A) Loss of lectins (Δ*lecA* and Δ*lecB*), quorum sensing (Δ*lasR*) and exopolysaccharide components (Δ*pel/psl*) did not change the aggregate assembly type and aggregates were assembled in stacked form similar to those seen in PAO1. (B) Stacked aggregates formed by cells lacking lectins (Δ*lecA* and Δ*lecB*), quorum sensing (Δ*lasR*) and exopolysaccharide components (Δ*pel/psl*) were the same size as PAO1 aggregates (Kruskal-Wallis, Dunn’s multiple comparison test, *p=*0.1, *p=*0.6, *p>*0.999 and *p>*0.999 when aggregate volumes of Δ*lecA*, Δ*lecB*, Δ*lasR* and Δ*pel/psl* were compared to PAO1; error bars are median with interquartile range of aggregate volume).

### Loss of OSA leads to clumped aggregates, independent of polymer and cell density

To determine differences in stacked and clumped aggregate assembly in SCFM2, despite the presence of polymers, we monitored aggregate assembly of PAO1 and PAO1Δ*wbpL* over time. We found that an increase in initial cell density resulted in the rapid formation of stacked aggregates, while there were no changes in clumped aggregate assembly (Movie S1 and S2).

When we monitored the aggregate formation of PAO1 and PAO1Δ*wbpL* over 6 h, we found that there was a significant change in PAO1 aggregate biovolume after 180 min of cells growing in SCFM2 and when the stacks were assembled (Fig. 4A) whereas, regardless of cell density, the biovolume of PAO1Δ*wbpL* aggregates remained constant over time (Fig. 4B). We also found that reducing the concentration of both polymers in SCFM2, led to the dissolution of stacked aggregates as expected, while it did not affect the formation of PAO1Δ*wbpL* clumped aggregates (Fig. S4). These findings suggest that loss of OSA prevents entropically-derived stacked aggregate assembly, and the associated mechanism is independent of the polymer concentration and/or cell density. This is a well-studied manifestation of aggregation of hydrophobic particles in colloidal environments (45, 46).

**Fig. 4.**
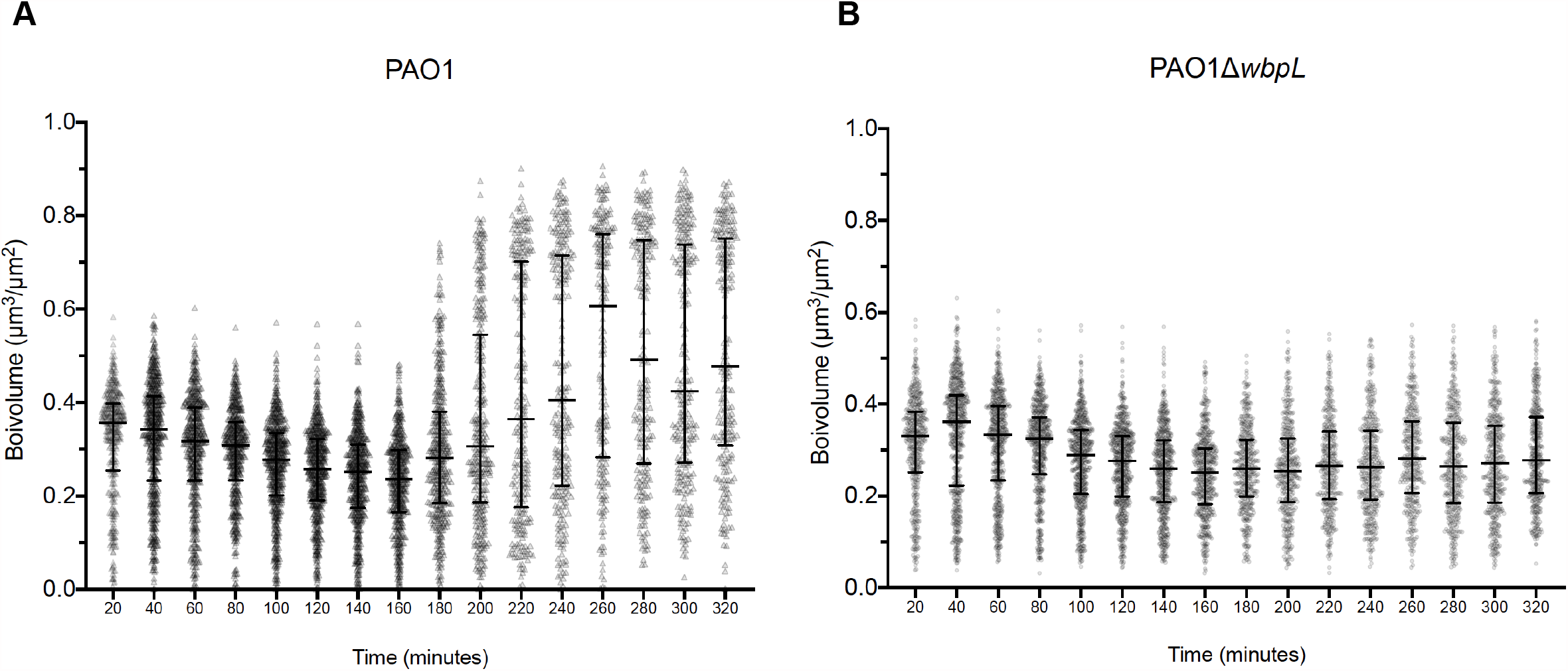
Clumped aggregate assembly is not dependent on cell density. (A) The aggregate biovolume of PAO1 significantly increased after 180 min of growth (median biovolume = 0.34-0.75 over time). (B) In PAO1Δ*wbpL* (lacking OSA), biovolume remained the same over time (median biovolume = 0.33-0.27 over time).

### Aggregate assembly of *P. aeruginosa* is not serotype specific but dependent on cell surface relative hydrophobicity

There are 20 serotypes of *P. aeruginosa* based on the glycosyl groups of OSA (38). As our findings were limited to PAO1 (serotype O5), we examined aggregate formation of PA14 (serotype O10), PAK (serotype O6) and STO1 (serotype O1) which all differ in the oligosaccharide units of OSA (38). In all serotypes we observed a stacked assembly similar to PAO1, but in an STO1 strain lacking OSA (Δ*wbpM*), we identified small clumped aggregates, with restoration of stacks when *wbpM* was complemented *in trans* (Fig. 5A & B). This data confirms that aggregate assembly type is not serotype specific.

**Fig. 5.**
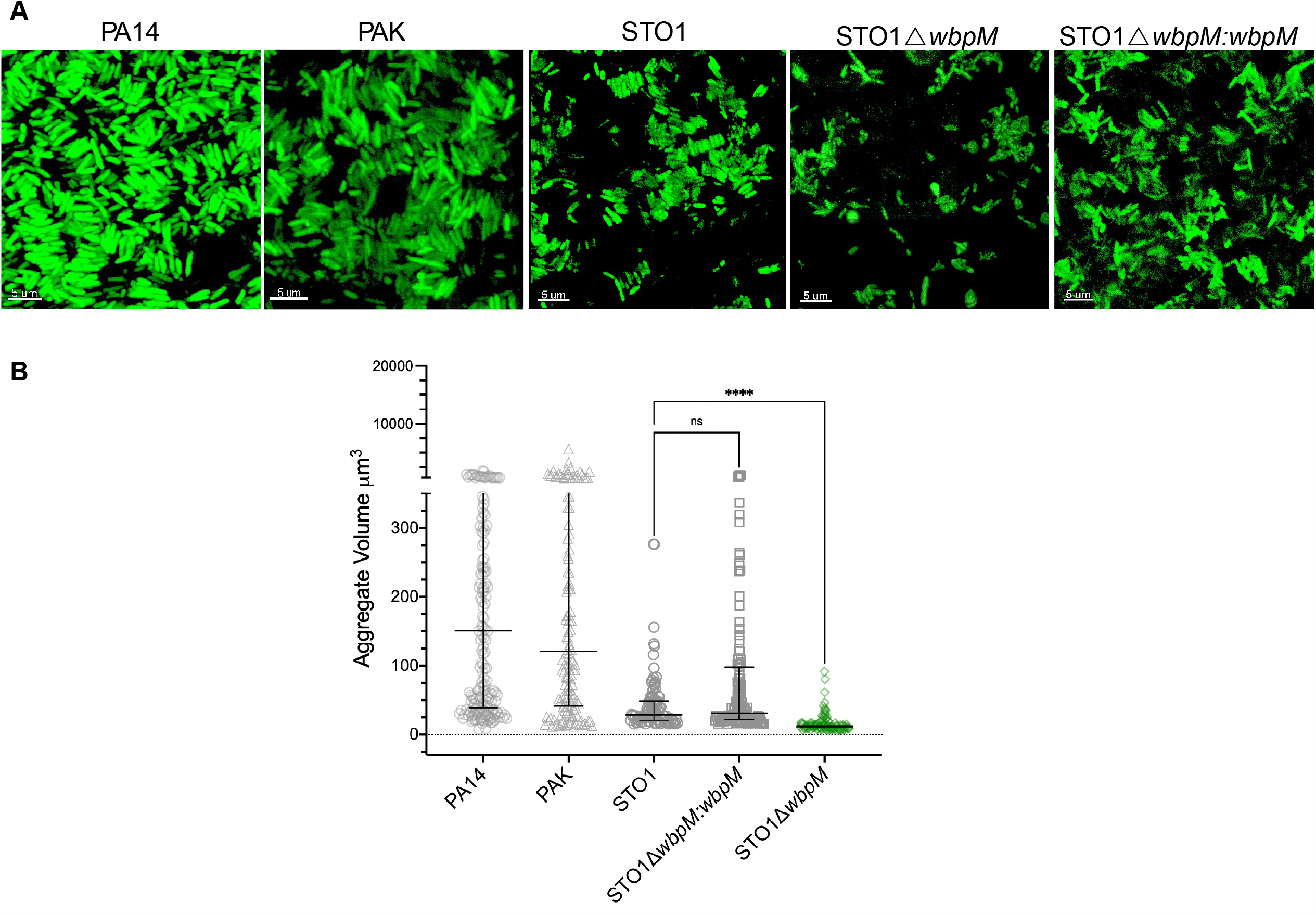
Clumped Aggregate assembly of *P. aeruginosa* is not serotype specific. (A) *P. aeruginosa* PA14 and PAK and STO1 formed stacked aggregates in SCFM2 and OSA^−^ mutant STO1 (STO1Δ*wbpM*) lead to clumped assembly of aggregates. (B) Loss of OSA altered aggregate assembly from stacked to clumped in STO1 and significantly decreased aggregate volume (Kruskal-Wallis, Dunn’s multiple comparison test, *p =* 0.0048; error bars are median with interquartile range of aggregate volume, each data point is representative of an aggregate).

Previously it has been shown that lack of OSA increases the hydrophobicity of the *P. aeruginosa* cell surface (30). To determine whether hydrophobicity correlated with aggregate type, we assessed the relative surface hydrophobicity of OSA+ and OSA-strains. We found a significant increase in surface hydrophobicity in OSA mutants (Fig. 6A), which corresponded with a clumping aggregate phenotype (Fig. 2). Loss of OSA is a common adaptive trait of *P. aeruginosa* in CF airways, possibly leading to increase in cell surface hydrophobicity that could alter spatial organization of *P. aeruginosa* cells in CF airways. We therefore evaluated the relative hydrophobicity of 11 *P. aeruginosa* isolates collected from the expectorated sputa of 2 individuals with CF. We observed heterogeneity in cell surface relative hydrophobicity of the CF isolates across the two patients (Fig. 6B).

**Fig. 6.**
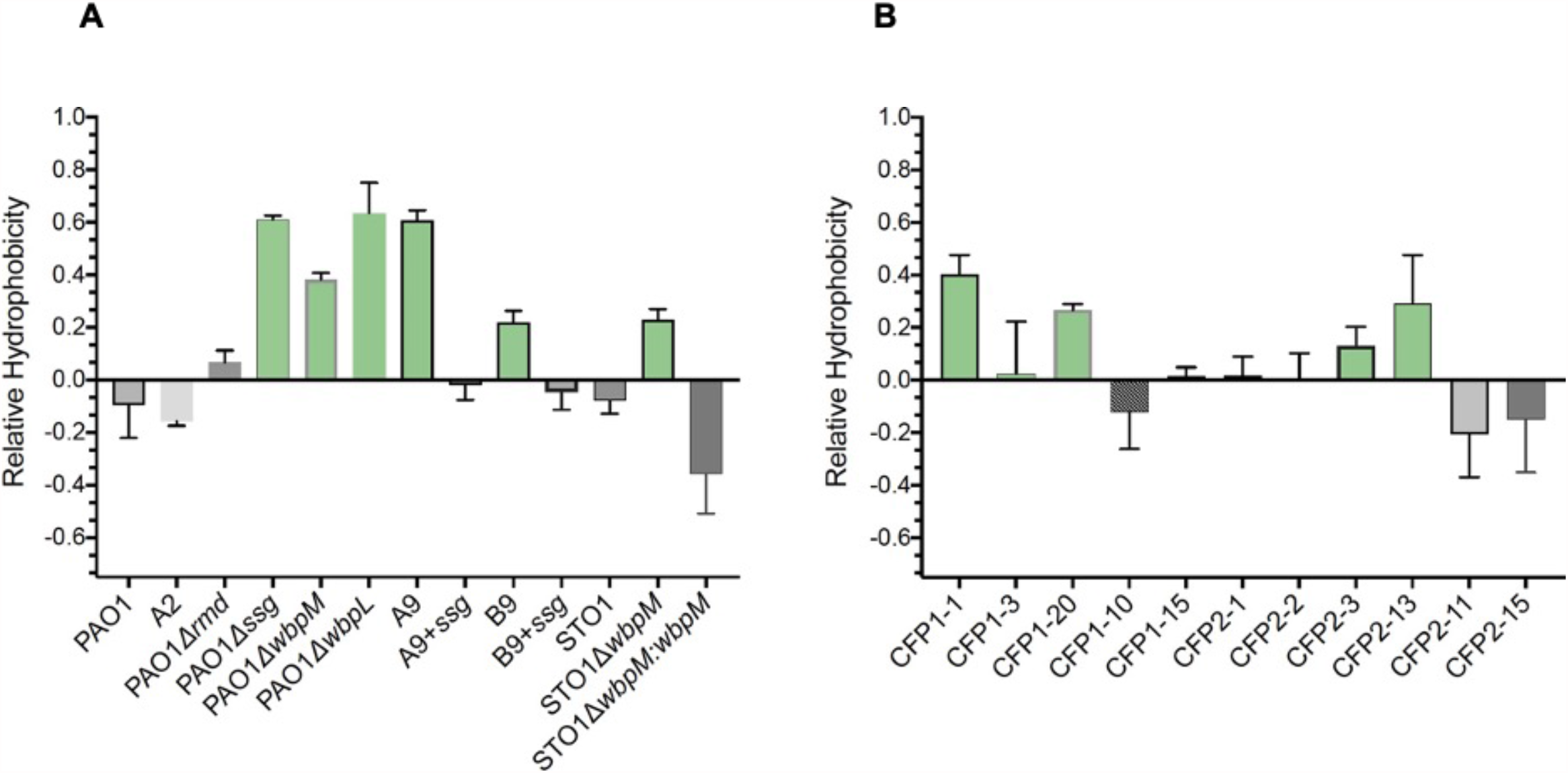
Cell surface hydrophobicity determines the aggregate assembly type. (A) The relative cell surface hydrophobicity was dependent on OSA, and mutation in *ssg, wbpL* and *wbpM* lead to an increase in relative hydrophobicity in PAO1 and STO1 (Green bars). (B) There was heterogeneity in the relative hydrophobicity of cell surfaces in *P. aeruginosa* isolates collected from two CF expectorated sputum samples (CFP1 and CFP2).

## Discussion

Despite a large body of work showing how *P. aeruginosa* adapts to the CF lung environment (19-21, 33, 47, 48), and population heterogeneity during chronic infection (20, 21, 23, 49, 50), little is known about the impact of this intra-specific heterogeneity on the formation of *P. aeruginosa* aggregates. To test whether genetic heterogeneity impacts aggregation, we investigated the aggregate formation of selected evolved isolates from a previous 50-day biofilm evolution experiment of PAO1 (23) in SCFM2 (24). We found that (i) there are two distinct types of aggregate assemblies formed by *P. aeruginosa* in SCFM2; (ii) the OSA impacts aggregate assembly type and (iii) the loss of OSA and LPS core+1 increases the hydrophobicity of the bacterial surface which prevents depletion aggregation.

Previously it was shown that aggregation of *P. aeruginosa* cells in a polymer rich environment can be due to depletion forces, where reduction of the free energy by increased entropy of the whole system leads to a stacked aggregation of bacterial cells (27). Change of the polymer electrostatic properties in the same study, altered the aggregate assembly to bridging assembly, suggesting that aggregate assembly type is driven by physical properties of the environment, and that the biological properties of the cells assumes little or no role on aggregate assembly type (27). However, we observed two distinct types of aggregate assembly by genetically diverse PAO1 isolates in SCFM2. Stacked aggregation is dependent on cell density and polymer concentration, and that increase in entropy is the driving force behind these assemblies. In contrast, clumping assembly is driven by changes in surface properties of *P. aeruginosa* cells. This indicates that although aggregate formation is influenced by physical forces, OSA strongly impacts upon the spatial organization enforced by physical properties of the environment.

Other studies have suggested that during CF infection, polymers like mucin can disperse cells in established biofilms (51), although our work suggests that polymers are more likely to influence the spatial arrangement of the cells. The clumping aggregate assembly was not influenced by factors that have previously been shown to be involved in biofilm and aggregate formation including lectins (42, 52), QS (6, 9, 53) and EPS (54-57). While we found that stacked aggregates had a larger biovolume than clumped aggregates, and this can be explained by depletion aggregation, other explanations also exist. First, stacked aggregates might be more likely to support growth within the aggregate leading to larger aggregates. Second, as clumped aggregates get larger, they may be more prone to disaggregation, leading to them never achieving a larger size even if there were cellular division occurring in the aggregate.

It remains to be determined whether stacked aggregates are found during human CF infection, although a recent study demonstrated that in CF airways, eDNA surrounds aggregates, suggesting that in CF lungs, eDNA could increase entropic force and form aggregates via depletion aggregation (58). It is noteworthy that stacked aggregates form in other species of bacteria such as in *Rhizobium leguminosarum* which was dependent on LPS core and O-antigen (59). Further, honeycomb structures of *P. aeruginosa* (multi-layers of stacks) were shown to from in the crops of *Drosophila melanogaster*, highlighting that stacked aggregates are found in some types of infection (60).

By assessing the relative hydrophobicity of cells forming different aggregate types, we found that *P. aeruginosa* cells that form clumped aggregates have higher relative hydrophobic surfaces. In agreement with previous studies (29, 61-63), we showed that this was due to a loss of OSA and exposure of uncapped LPS core. Regulating aggregate assembly type by directly altering OSA expression levels in environments containing differential levels of cells and polymers (such as CF sputum), may allow cells to better resist environmental stressors such as the host immune response, antibiotics or phage. The loss of OSA has been reported in *P. aeruginosa* strains isolated from CF sputum, suggesting that finding small, clumped aggregates in CF sputum and CF airways (64), might be directly due to the loss of OSA. As our work was performed *in vitro*, further studies are required to determine how OSA and surface hydrophobicity impacts on aggregate formation in *P. aeruginosa* isolates growing in CF lungs.

Overall, our findings highlight that the surface properties of *P. aeruginosa* cells determines how they form aggregates in environments with different physicochemical properties, providing potential benefits from social interactions with highly-related cells. Our findings also highlight, that changes in cell surface properties may influence how aggregates form in other species of bacteria and provides explanations as to how different *P. aeruginosa* strains and species can stably co-exist in microbiomes.

## Materials & Methods

### Bacterial strains and culture condition

We selected 7 evolved isolates of PAO1 from 50 day evolved populations in SCFM (23). We transformed all *P. aeruginosa* strains used in this study with pME6032:*gfp* plasmid (65) using electroporation (66). Briefly, to prepare electrocompetent *P. aeruginosa* cells; we grew the bacterial cells in LB broth overnight, we then washed the overnight cultures with 300 mM sucrose solution at room temperature, and then resuspended the bacterial pellets in 1ml of 300 mM sucrose. We then electroporated 50 µl of electrocompetent cells with 2 µl of purified plasmid and recovered the cells by addition of 950 µl of LB broth and incubation at 37 °C/ 200 rpm for 30 mins. We selected the transformed cells by plating out the electroporated bacteria onto LB agar plates supplemented with 200 µg/ml of tetracycline. We obtained the clinical isolates from Emory CF@LANTA Research Center. Patients in this study were aged between 21-29 at the time of collection of the sputum samples. This study was approved by Institutional Review Board (IRB) at Georgia Institute of Technology and Emory University Hospital. A list of all bacterial strains used in this study is available in Table S2.

### Determining diversity in colony morphologies

To determine diversity in colony morphology in evolved populations (23), we used a Congo Red based agar media (1% agar, 1×M63 salts (3g monobasic KHPO_4_, 7 g K_2_ PO_4_, 2 g NH_4._ 2SO_4_, pH adjusted to 7.4), 2.5 mM magnesium chloride, 0.4 mM calcium chloride, 0.1% casamino acids, 0.1% yeast extracts, 40 mg/l Congo red solution, 100 µM ferrous ammonium sulphate and 0.4% glycerol) (67). We inoculated each evolved isolate in LB broth and incubated for 6 h at 37°C/200 rpm, then we spotted a 10 µl of the culture onto Congo Red agar plates. We incubated the plated at 37°C for 24 h and a further 4 days at room temperature.

### Genomic DNA extraction and whole genome sequencing

We plated each of the selected evolved isolates on LB agar plates, picked single colonies of each isolate and then inoculated in 5 ml of SCFM (35) and incubated overnight at 37 °C/ 200 rpm. We extracted the genomic DNA using the QIAGEN Blood and Tissue DNAeasy kit. We prepared sequencing libraries using the NexteraXT protocol (Illumina), and sequenced in 24-plex on the Illumina MiSeq platform to obtain an approximate calculated level of coverage of 50× for each evolved isolate. For SNP calling, we used breseq analysis (consensus mode) (37, 68, 69) and compared the genetic variation in each evolved isolate to the PAO1 ancestral strain. The sequences are available at the NCBI SRA database (PRJNA702741).

### Image acquisition and analysis

For imaging aggregates in SCFM2 (24), we inoculated each bacterial isolate into TSB-broth supplemented with 200 µg/ml of tetracycline and incubated at 37 °C/200 rpm overnight. We inoculated 50 µl of the overnight culture into 5 ml of SCFM and incubated at 37 °C/ 200 rpm for 5-6 h, until cultures reached mid-log phase (OD_600_ = 0.5). We then adjusted the OD_600_ to 0.05 in 400 µl of freshly made SCFM2 containing 0.6 mg/ml of DNA and 5 mg/ml of mucin (8, 24). We incubated the cultures at 37 °C for 16 h in chamber slides (Lab-Tek®) before image acquisition. We used a confocal LSM880 microscope equipped with a 63× oil immersion lens for image acquisition and scanned the aggregates using diode laser 488 nm, and collected fluorescent emission between 480-530 nm for image acquisition. For imaging the cells grown in SCFM, we adjusted the OD_600_ of cells from mid-log phase growth to 0.05 in 400 µl of freshly made SCFM. We incubated the cultures at 37 °C for 16 h in chamber slides before image acquisition. For image analysis, we used Imaris 9.0.1 image analysis software to analyze the morphological properties of the aggregates and measured the surface area and volume of each aggregate using a surface model algorithm. We used the same parameters for particle and voxel size. We measured the aggregate volume and surface area in 10 images acquired for each strain in three independent experiments (over 1000 aggregates were measured in total for each condition). For time course experiments, we used the same image acquisition parameters, using the time series option and imaged as Z stacks every 20 min for up to 10 h. To assess the role of bacterial cell density on aggregation, we adjusted the OD_600_ to 0.1 in 400 µl of SCFM2 and imaged the cells every 20 min for 6 h. We prepared time series videos using the 3D plugin in Fiji (70) and Adobe Lightroom.

### Gene deletion and complementation

We used standard genetic techniques for the construction of *P. aeruginosa* mutants. To delete *ssg, rmd, wbpL, wbpM, waaL, wzy, wzz1* and *wzz2*, we PCR amplified 600 bp DNA sequences flanking the open reading frame of each gene using Q5 DNA polymerase (New England Biolabs). We then cloned these sequences into EcoRI-XbaI digested pEXG2 by Gibson assembly using NEBuilder HiFi Assembly master mix (New England Biolabs) and transformed into *E*.*coli* S17 λpir. We verified cloned inserts by colony PCR and Sanger sequencing (Eurofins Genomics). We introduced the deletion constructs into PAO1 by electroporation and selected strains carrying single crossover insertions of the deletion constructs on LB agar plates supplemented with 100 µg/ml gentamycin. We cultured gentamycin resistant colonies in LB without antibiotic and plated on LB agar plates with 0.25% NaCl and 5% sucrose. We then selected sucrose resistant colonies and screened them for gentamycin sensitivity to ensure loss of the pEXG2 construct and assessed them for the desired gene deletion by colony PCR and Sanger sequencing of the PCR product. For *ssg* complementation we PCR amplified the *ssg* coding sequence and 100 bp upstream sequence (including the *ssg* native promoter) using Q5 DNA polymerase (New England Biolabs). We cloned this 1057 bp product into KpnI-BamHI digested pUC18T-miniTn7-Gent by Gibson assembly using NEBuilder HiFi Assembly master mix (New England Biolabs) and transformed into *E*.*coli* S17 λpir. We verified the cloned insert by colony PCR and Sanger sequencing (Eurofins Genomics). We co-transformed the complementation construct with the Tn7 helper plasmid pTNS3 into PAO1Δ*ssg* and evolved isolates by electroporation and selected on LB agar plates supplemented with 100 µg/ml gentamycin. We verified the strains for *ssg*^+^ complementation by colony PCR and for loss of pUC18-miniTn7-Gent vector and pTNS3 by screening for carbenicillin sensitivity.

### LPS extraction

We isolated bacterial lipopolysaccharide by the hot phenol extraction method (71). Briefly, we pelleted 5 ml overnight cultures of PAO1 and PAO1-derived strains in LB broth by centrifugation for 10 min at 4200× *g*. We resuspended the pellets in 200 µl 1× SDS buffer (2% β-mercaptoethanol (BME), 2% SDS, 10% glycerol, 50 mM Tris-HCl, pH 6.8) and incubated at 99°C for 15 min. Then we added 5 µl of 20 mg/ml proteinase K (Sigma) to each tube and incubated cell lysates at 59°C for 3 h. Then we added 200 µl of ice-cold Tris-saturated phenol to each sample, vortexed for 10 min, then added 1 ml diethyl-ether and vortexed for a further 10 sec. We centrifuged the samples for 10 min at 16000× *g* and extracted the bottom layer. We performed a second extraction with phenol and diethyl-ether as above. We mixed an equal volume of the extracted LPS samples with an equal volume of 2× SDS buffer and electrophoresed 10 µl of each sample on Novex 4-20% plyacrylamide gradient gels (ThermoFisher) in Tris-Glycine-SDS buffer. Following electrophoresis, we visualized LPS with a ProQ Emerald Lipopolysaccharide staining kit (ThermoFisher).

### Assessing cell surface hydrophobicity

To assess the levels of cell surface hydrophobicity, we used hydrophobic interaction chromatography (29). Briefly, we grew bacterial cells for 6-8 h at 37 °C/ 200 rpm, to reach mid-log phase. We harvested the cells and washed the cells 3× with ice cold 3M NaCl pH=7 and resuspended in 3M NaCl. We used octyl-Sepharose CL-4C beads (SIGMA) to assess the interaction of hydrophobic cells to these beads compared to control Sepharose CL-4C beads (SIGMA). We prepared bead columns by a 3× wash of the beads with Mili-Q water, and then 3× washes with 3M NaCl (pH=7) (at 4° C). We then prepared 1 ml columns of both beads by using 3 ml diameter filter paper. We added 100 µl of bacterial suspension and incubated at room temperature for 15 min. We measured the OD_450_ of flow through from each column. We calculated the relative hydrophobicity based on the ratio of OD_450_ octyl-Sepharose CL-C4 column flow through and control column.

### Statistical analysis

For statistical analysis of the aggregate volume and biovolume distribution, we used GraphPad Prism 8.0.

## Supplemental figure legends

**Fig. S1.**
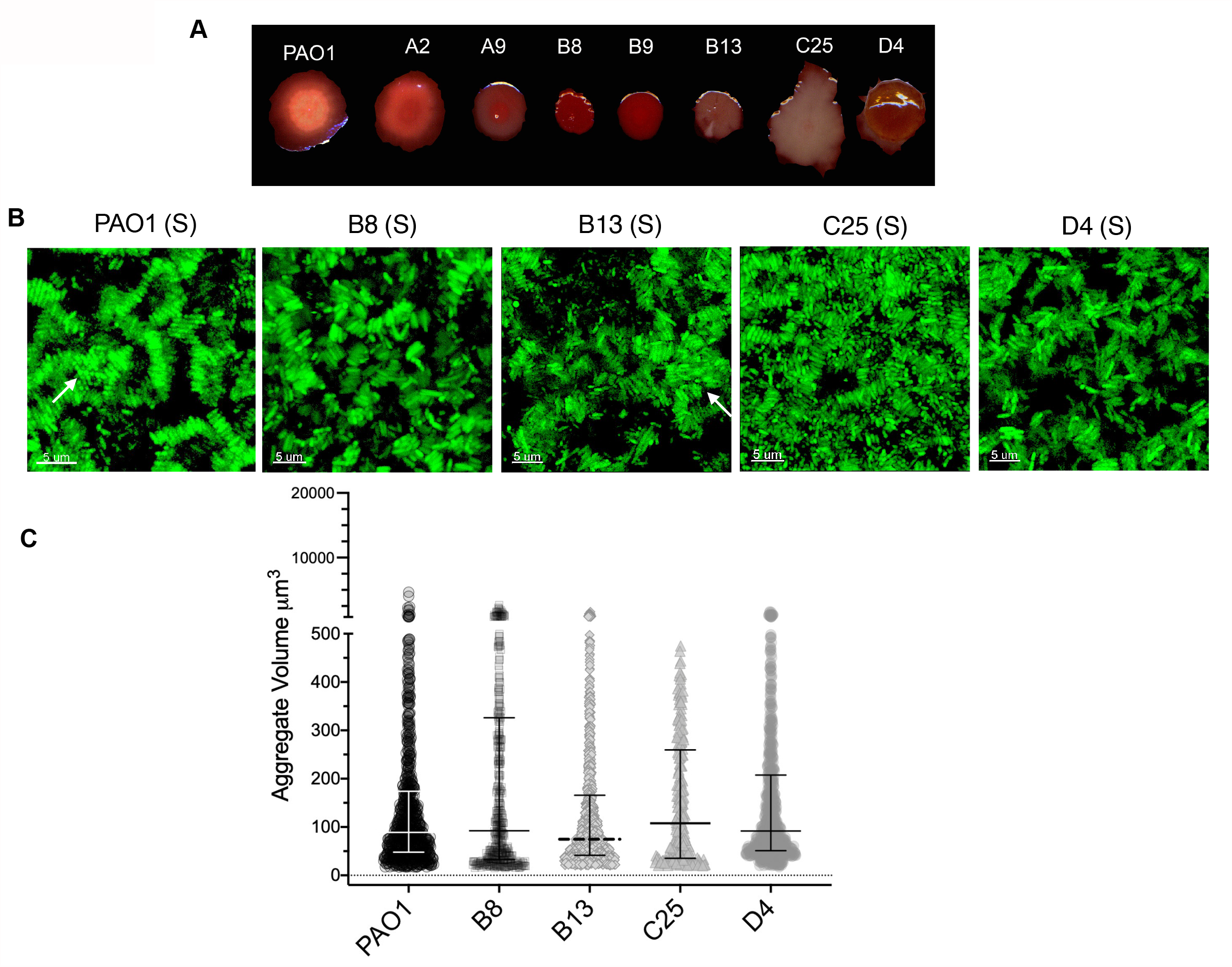
Aggregate assembly of PAO1 evolved isolates in SCFM2. (A) Evolved isolates of PAO1 displayed differential colony morphologies on Congo red agar plates. (B) Evolved isolates B8, B13, C25 and D4 displayed stacked aggregate assembly (labeled as S). (C) There were no significant differences in the volume of stacked aggregates in these isolates compared to PAO1 (*p <*0.0001, Kruskal-Wallis, Dunn’s multiple comparison test, error bars are median with interquartile range of aggregate volume).

**Fig. S2.**
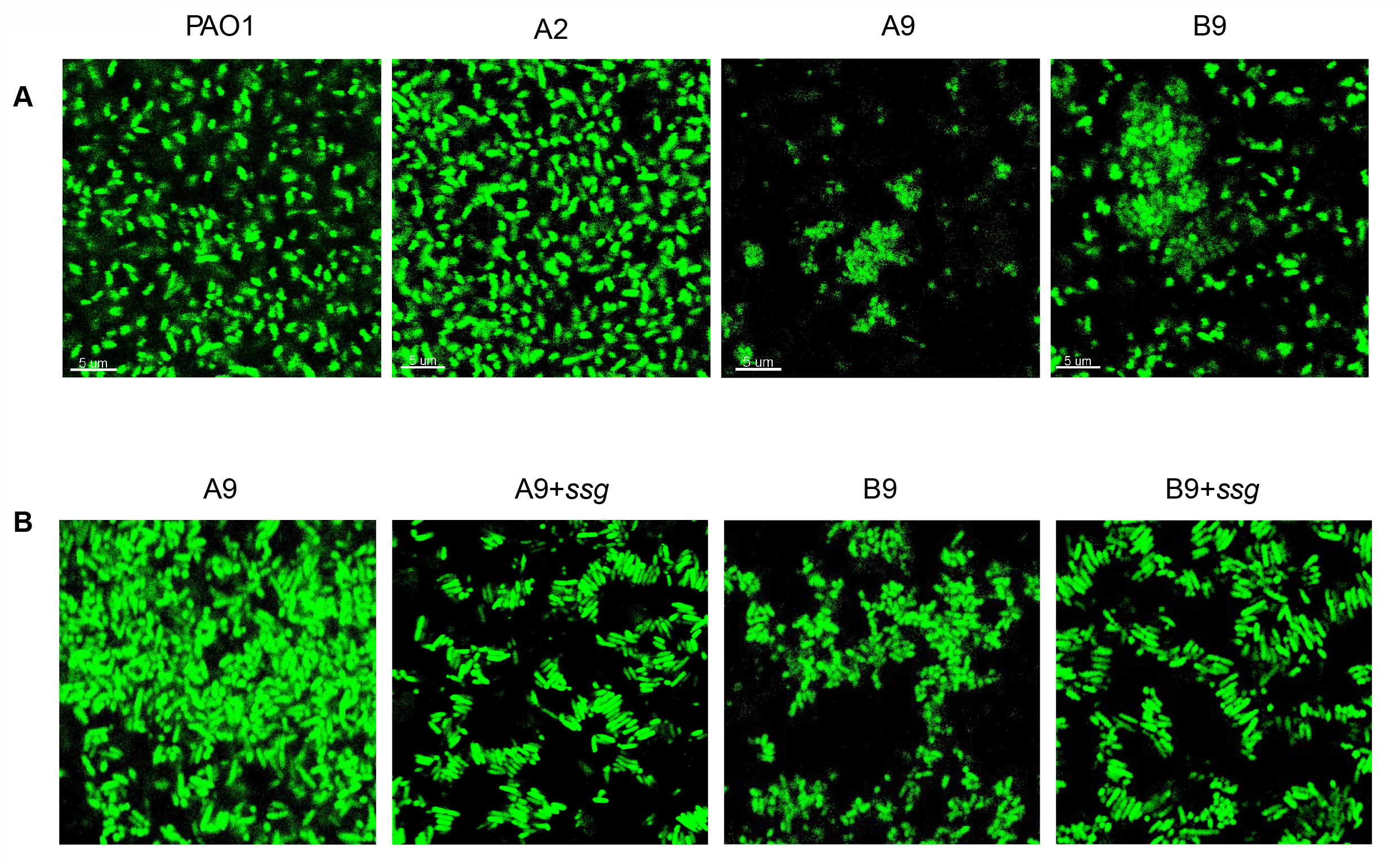
Mutation in PA5001 (*ssg*) in PAO1 switches the stacked aggregate assembly to clumped assembly. (A) PAO1 and A2 cells formed a dispersed layer of cells on cover slips when grown in SCFM (without any polymer), while A9 and B9 formed small clumps in SCFM. (B) Complementing A9 and B9 isolates with *ssg* restored the aggregate assembly to stacked (Kruskal-Wallis, Dunn’s multiple comparison test, *p*<0.0001, error bars are median with interquartile range of aggregate volume).

**Fig. S3.**
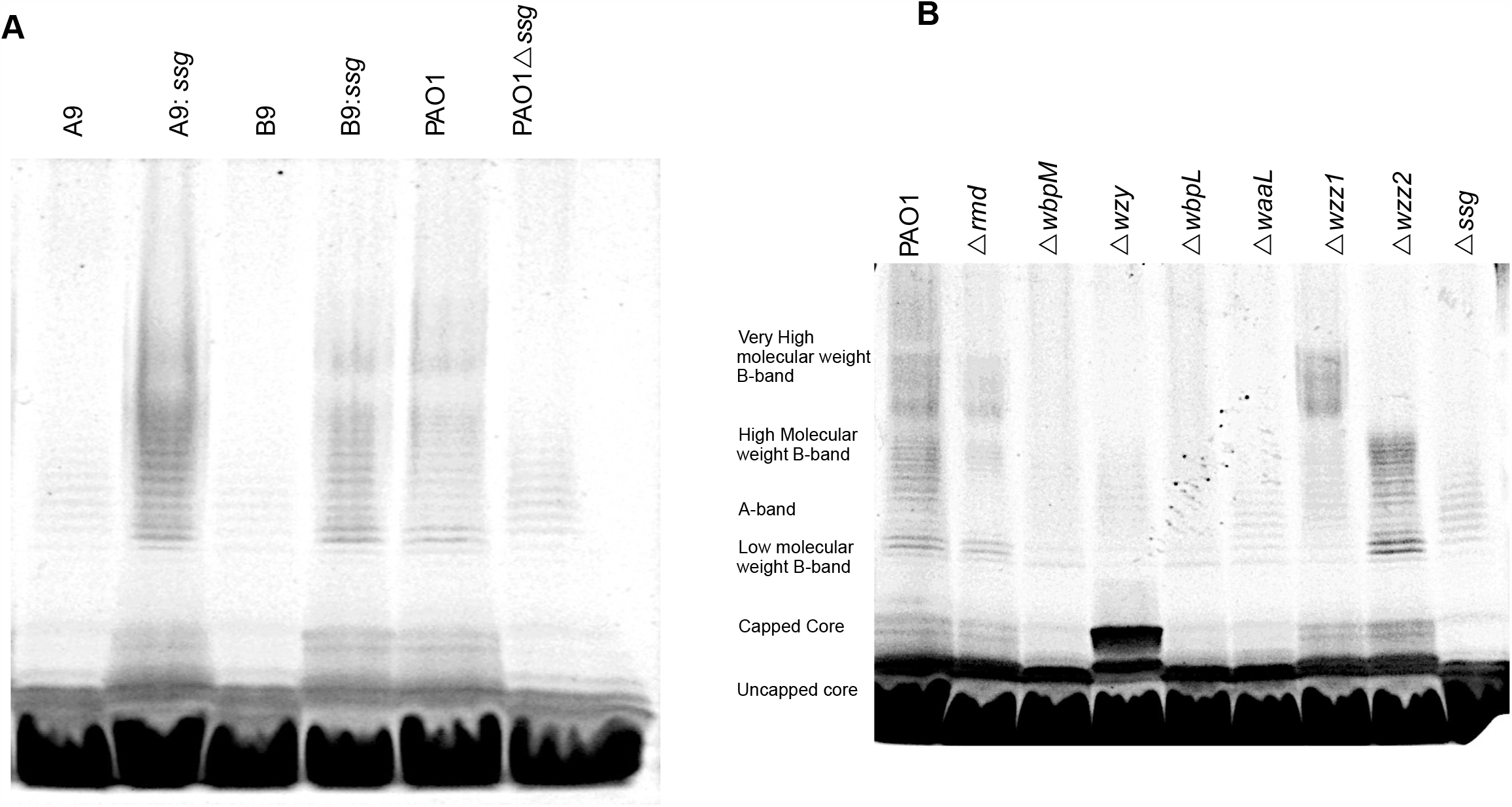
Loss of *ssg* function changes the LPS OSA profile in *P. aeruginosa*. (A) Evolved isolates of PAO1 with 1 bp deletion in *ssg* had the same OSA profile as PAO1Δ*ssg*, and complementation of *ssg* in *trans* restored the OSA pattern. (B) Loss of *ssg* resulted in the loss of OSA and a change in capped core pattern.

**Fig. S4.**
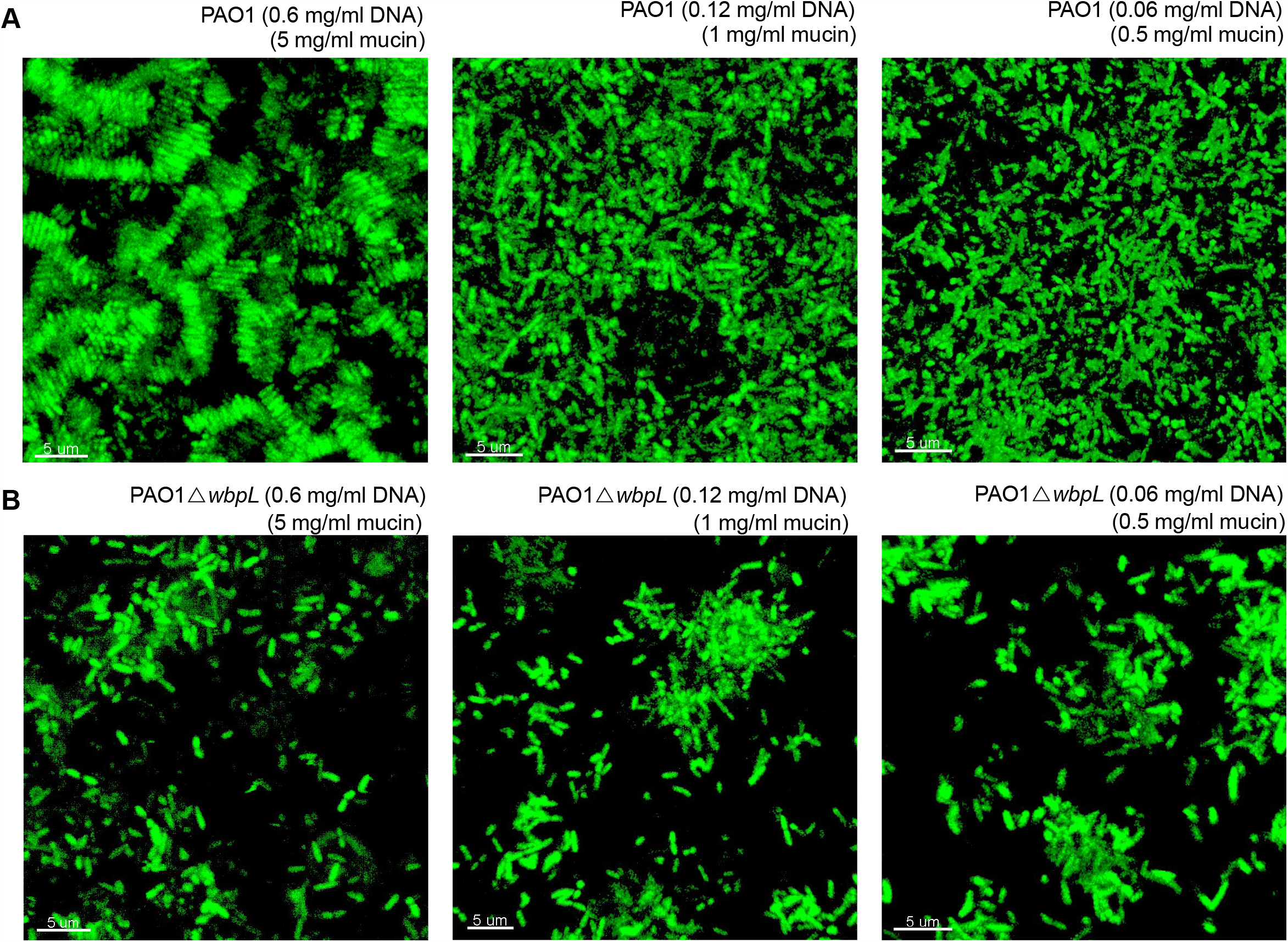
Stacked aggregate assembly is dependent on the concentration of both eDNA and mucin. (A) Stacked aggregate formation was disrupted by diluting polymers in SCFM2. (B) Clumped aggregate assembly was independent of the polymer concentration in the environment.

**Movie S1. Increased cell density reduced the time of stacked aggregation**. To determine whether stacked aggregate assembly was dependent on cell density, we monitored the growth of PAO1 over time (initial density adjusted OD_600_ to 0.1 in 400 µl of SCFM2). Stacked aggregates were assembled after 180 min of growth.

**Movie S2. Increased cell density does not impact the timing of formation of clumped aggregation**. We monitored the growth of PAO1Δ*wbpL* (initial density adjusted OD_600_ to 0.1 in 400 µl of SCFM2). There was no change in aggregate assembly over time.

**Dataset 1. List of SNPs in 7 evolved isolates of PAO1**.

**Table S1. Stacked aggregates have higher biovolume than clumped aggregates**. To determine the differences in biomass of each aggregate type, we calculated the total biovolume (µm^3^ /µm^2^) of all aggregates in each acquired image, using Imaris. There was a significant difference between distribution of biovolume of stacked and clumped (green) aggregates (Kruskal-Wallis, Dunn’s multiple comparison test, *p*<0.0001; error bars are median with interquartile range of aggregate biovolume).

**Table S2. List of the strains used in this study**.

## Acknowledgments

For funding, we thank the Georgia Institute of Technology; The Cystic Fibrosis Foundation for grants (DIGGLE18I0; DIGGLE20G0) to SPD and a Fellowship to SA (AZIMI18F0); CF@LANTA for a Fellowship to SA (3206AXB); The National Institute for Health (R01AI153116) to SPD; the National Science Foundation (1806606) to JEC. We thank Marvin Whiteley for useful discussion and use of his Confocal Ziess LSM880 and Imaris image analysis platform. Access to the CF Biospecimen Registry (CFBR) at Emory Children’s Center for Cystic Fibrosis and Airways Disease Research was provided through Children’s Healthcare of Atlanta and the Emory University Pediatric CF Discovery Core. We thank Arlene Stecenko and Katy Clemmer for assistance acquiring bacterial isolates.

