## Supplementary material for "O-specific antigen-dependent surface hydrophobicity mediates aggregate assembly type in *Pseudomonas aeruginosa*": Table S1

| | PAO1 | A2 | A9 | B9 |
|---|---|---|---|---|
| Minimum | 0.110031 | 0.12282713 | 0.100064 | 0.10049691 |
| 25% Percentile | 0.248399 | 0.267627 | 0.222612 | 0.15592075 |
| Median | 0.35255 | 0.312206 | 0.277916 | 0.2315546 |
| 75% Percentile | 0.41007625 | 0.393158 | 0.33127 | 0.33155867 |
| Maximum | 0.559253 | 0.600513 | 0.371328 | 0.469442558 |
