## Supplementary material for "O-specific antigen-dependent surface hydrophobicity mediates aggregate assembly type in *Pseudomonas aeruginosa*": Table S2

| *P. aeruginosa* strain | Description | | Source |
|---|---|---|---|
| **PAO1** | Wild type PAO1 (Nottingham strain) | | Holloway collection |
| **A2** | Evolved PAO1 isolate | Evolved isolate | This study |
| **A9** | Evolved PAO1 isolate | Evolved isolate | This study |
| **A9:*ssg*** | Evolved PAO1 isolate complemented with *ssg* gene | Evolved isolate | This study |
| **B8** | Evolved PAO1 isolate | Evolved isolate | (1) |
| **B9** | Evolved PAO1 isolate | Evolved isolate | This study |
| **B9:*ssg*** | Evolved PAO1 isolate complemented with *ssg* gene | Evolved isolate | This study |
| **B13** | Evolved PAO1 isolate | Evolved isolate | This study |
| **C25** | Evolved PAO1 isolate | Evolved isolate | This study |
| **D4** | Evolved PAO1 isolate | Evolved isolate | This study |
| **PAO1Δ*ssg*** | PAO1 with *ssg* deletion | Isogenic mutant | This study |
| **PAO1Δ*rmd*** | PAO1 with *rmd* deletion/CPA- | | This study |
| **PAO1Δ*wbpM*** | PAO1 with *wbpM* deletion/OSA- | | This study |
| **PAO1Δ*wbpL*** | PAO1 with *wbpL* deletion/ CPA- OSA- | | This study |
| **PAO1Δ*waal*** | PAO1 with *waal* deletion/ OSA- | | This study |
| **PAO1Δ*wzy*** | PAO1 with *wzy* deletion/ OSA- | | This study |
| **PAO1Δ*wzz1*** | PAO1 with *wzz1* deletion/No high molecular weight B-band | | This study |
| **PAO1Δ*wzz2*** | PAO1 with *wzz2* deletion/No very high molecular weight B-band | | This study |
| **STO1** | Clinical isolate of *P. aeruginosa* serotype O-1 | | [2] |
| **PA14** | PA14 wildtype | | |
| **PAK** | PAK wildtype | | |
| **STO1Δ*wbpM*** | Clinical isolate of *P. aeruginosa* serotype O-1/ OSA- | | [2] |
| **STO1Δ*wbpM:wbpM*** | Clinical isolate of *P. aeruginosa* serotype O-1; OSA- / complemented with serotype O-1 *wbpM* | | [2] |
| **PAO1Δ*lecA*** | PAO1 with *lecA* deletion | | [3] |
| **PAO1Δ*lecB*** | PAO1 with *lecB* deletion | | This study |
| **PAO1Δ*pel/psl*** | PAO1 with *pel/psl* deletions | | [4] |

| | | | |
|---|---|---|---|
| **PAO1Δ*lasR*** | PAO1 with *lasR* deletion | | This study |
| **CFP1-1** | CF isolate of *P. aeruginosa, isolated from* CFP1 sputum sample | Clinical isolate | This study |
| **CFP1-3** | CF isolate of *P. aeruginosa, isolated from* CFP1 sputum sample | Clinical isolate | This study |
| **CFP1-10** | CF isolate of *P. aeruginosa, isolated from* CFP1 sputum sample | Clinical isolate | This study |
| **CFP1-15** | CF isolate of *P. aeruginosa, isolated from* CFP1 sputum sample | Clinical isolate | This study |
| **CFP1-20** | CF isolate of *P. aeruginosa, isolated from* CFP1 sputum sample | Clinical isolate | This study |
| **CFP2-1** | CF isolate of *P. aeruginosa, isolated from* CFP2 sputum sample | Clinical isolate | This study |
| **CFP2-2** | CF isolate of *P. aeruginosa, isolated from* CFP2 sputum sample | Clinical isolate | This study |
| **CFP2-3** | CF isolate of *P. aeruginosa, isolated from* CFP2 sputum sample | Clinical isolate | This study |
| **CFP2-11** | CF isolate of *P. aeruginosa, isolated from* CFP2 sputum sample | Clinical isolate | This study |
| **CFP2-13** | CF isolate of *P. aeruginosa, isolated from* CFP2 sputum sample | Clinical isolate | This study |
| **CFP2-15** | CF isolate of *P. aeruginosa, isolated from* CFP2 sputum sample | Clinical isolate | This study |


1. Azimi, S., et al., *Allelic polymorphism shapes community function in evolving Pseudomonas aeruginosa populations.* ISME J, 2020.
2. Davis, M.R., Jr., et al., *Identification of the mutation responsible for the temperature-sensitive lipopolysaccharide O-antigen defect in the Pseudomonas aeruginosa cystic fibrosis isolate 2192*. J Bacteriol, 2013. **195**(7): p. 1504-14.
3. Diggle, S. P., et al., *The galactophilic lectin, LecA, contributes to biofilm development in Pseudomonas aeruginosa*. Environ Microbiol, 2006. **8**(6): p. 1095-1104.
4. Irie, Y., et al., *The Pseudomonas aeruginosa PSL Polysaccharide Is a Social but Noncheatable Trait in Biofilms.* mBio, 2017. **8**(3).
